# Human personality reflects spatio-temporal and time-frequency EEG structure

**DOI:** 10.1101/317032

**Authors:** Anastasia E. Runnova, Vladimir A. Maksimenko, Maksim O. Zhuravlev, Pavel Protasov, Roman Kulanin, Marina V. Khramova, Alexander N. Pisarchik, Alexander E. Khramov

**Author notes:** These authors contributed equally to this work (Anastasia E. Runnova, Vladimir A. Maksimenko, Maksim O. Zhuravlev, Alexander E. Khramov).

## Abstract

The brain controls all physiological processes in the organism and regulates its interaction with the external environment. The way the brain solves mental tasks is determined by individual human features, which are reflected in neuronal network dynamics, and therefore can be detected in neurophysiological data. Every human action is associated with a unique brain activity (motor-related, cognitive, etc.) represented by a specific oscillatory pattern in a multichannel electroencephalogram (EEG). The connection between neurophysiological processes and personal mental characteristics is manifested when using simple psycho-diagnostic tests (Schulte tables) in order to study the attention span. The analysis of spatio-temporal and time-frequency structures of the multichannel EEG using the Schulte tables allows us to divide subjects into three groups depending on their neural activity. The personality multi-factor profile of every participant can be individually described based on both the Sixteen Personality Factor Questionnaire (16PF) and a personal interview with an experienced psychologist. The correlation of the EEG-based personality classification with individual multi-factor profiles provides a possibility to identify human personality by analyzing electrical brain activity. The obtained results are of great interest for testing human personality and creating automatized intelligent programs that employ simple tests and EEG measurements for an objective estimation of human personality features.

## Introduction

Every human activity involves a generation of particular patterns in electroencephalographic (EEG) recordings with common properties for different subjects. For instance, the perception of visual stimuli is known to induce an event-related response of the neuronal brain network, in particular, a decrease in alpha-wave (8–12 Hz) and an increase in beta-wave (15–30 Hz) activities. Such a behavior reflects different cognitive functions, namely, the alpha-wave suppression is associated with visual [1] or auditory [2] attention, while the beta-wave activation relates to information processing [3] and an alerted state [4,5]. Similar cognitive activity was observed in the group of motivation-dependent volunteers [6].

Universal EEG patterns were also identified in patients with pathological brain activity, e.g., epileptic seizures. This type of activity is closely related to global synchronization in the brain’s neuronal network [7], which is manifested as high-amplitude oscillations with a specific waveform (series of well-pronounced spikes and waves) and frequency [8]. Such spike-and-wave oscillations have similar properties for different patients and are only determined by the type of epilepsy.

Different physiological and psychological states (e.g., sleep stages, arousal, etc.) are known to possess specific properties of neural activity. For instance, motor-related brain activity is manifested in the brain as a specific scenario of neural activity with well-defined frequency and spatial localization. Particularly, it is characterized by event related desynchronization (ERD) in alpha/mu-and beta-bands [9]. The same features are observed during motor imagery in specially trained subjects [10, 11]. However, different scenarios occur in untrained subjects, where EEG patterns can vary from subject to subject [12]. Such a variation is caused by the task complexity when each subject chooses his own strategy to process the task, that results in individual time-frequency and spatio-temporal EEG structures. Along with motor imagery, the personality is more pronounced during mental task processing. It was also shown that human personality causes individual scenarios during decision-making [13] and affects learning performance [14].

We suppose that individual features of human personality, when we wish to define the ways of how a human processes mental tasks, affect neural network dynamics and therefore can be seen in EEG recordings. It should be noted that this problem was attacked yet in 1973. By analyzing resting states, Edwards and Abbott [15] tried to reveal personality traits in EEG signals. However, their attempt was unsuccessful because personality is not manifested when a person is at rest. Until now, this problem remains open [16]. In this context, the research focused on the assessment of personality based on the analysis of EEG data in the resting state has not reached consistent conclusions. While some papers reported a successful assessment [17], others concluded that resting state features could not be used [18]. Based on the previous studies, we hypothesize that the features associated with personality traits are more pronounced during cognitive activity. A similar assumption was made by Fink and Neubauer [19], who studied extraversion personality of subjects by dividing them into two groups according to a psychological test.

In the present work, we record multichannel EEG during the completion of mental tasks to reveal individual features of the brain activity related to personality. In order to verify our hypothesis, at the first stage, we analyze spatio-temporal and time-frequency EEG structures of subjects who performed the Schulte table test, in order to classify them according to their neural activity scenario. At the second stage, the personality multi-factor profile is created for every participant on the base of both the Sixteen Personality Factor Questionnaire (16PF) [20, 21] and a personal interview with an experienced psychologist. Finally, we compare the results of two classifications.

## Materials and methods

### Participants

Twenty two conditionally healthy men (33 ± 7 years), right-handed, amateur practitioners of physical exercises, and non-smokers participated at the experiment. All of them were asked to maintain a healthy life regime with an 8-hrs night rest during 48 hrs prior the experiment. All volunteers provided informed written consent before participating in the experiment. The experimental procedure was performed in accordance with the Helsinki’s Declaration and approved by the local Ethics Committee of the Yuri Gagarin State Technical University of Saratov.

### Experimental design

The experiments were carried out with each subject independently. The participants were not previously informed about the experiment conditions. The experimental research was conducted by independent researchers of various specializations and included two separate stages for each volunteer.

**The first experimental stage** was based on accepted techniques for the definition of a person’s psychological type. For every participant, a personality multi-factor profile was described on the base of both the Sixteen Personality Factor Questionnaire (16PF) [20, 21] and a personal interview with an experienced psychologist. The 16PF contained 185 items organized into 16 primary factor scales and was adapted for Russian language and cultural context features [22–26]. We used the fully automated version of the 16PF, i.e., no paper-and-pencil materials were used. In this automated version, the items appeared on the screen one by one. There was the option to return to the immediately preceding item to correct inadvertent keying errors. However, the participant was not able to browse through the items. The program saved raw scale scores for every test and item responses.

**The second stage of experimental work** was carried out during the first half of the day at a specially equipped laboratory where the volunteer was sitting comfortably. The influence of external stimuli, such as extraneous sounds and bright light, was minimized as much as possible. All participants performed a series of simple psycho-diagnostic tests using the Schulte tables [27–30]) to study their attention features (see Fig. 1 (a)), under direct supervision of a professional psychologist. The Schulte table is a 5 × 5 matrix of random numbers from 1 to 25, as shown in Fig. 1(b). The psychological task was to find all numbers in a reverse order. During these *active* experimental phases, each person had to complete *R* = 5 tables. For every *i* testing series, the completion time *T_i_* was registered. Between the active phases, each volunteer had a short resting interval referred to as a *passive* experimental phase. The experimental design is shown in Fig. 1 (c).

**Fig 1.**
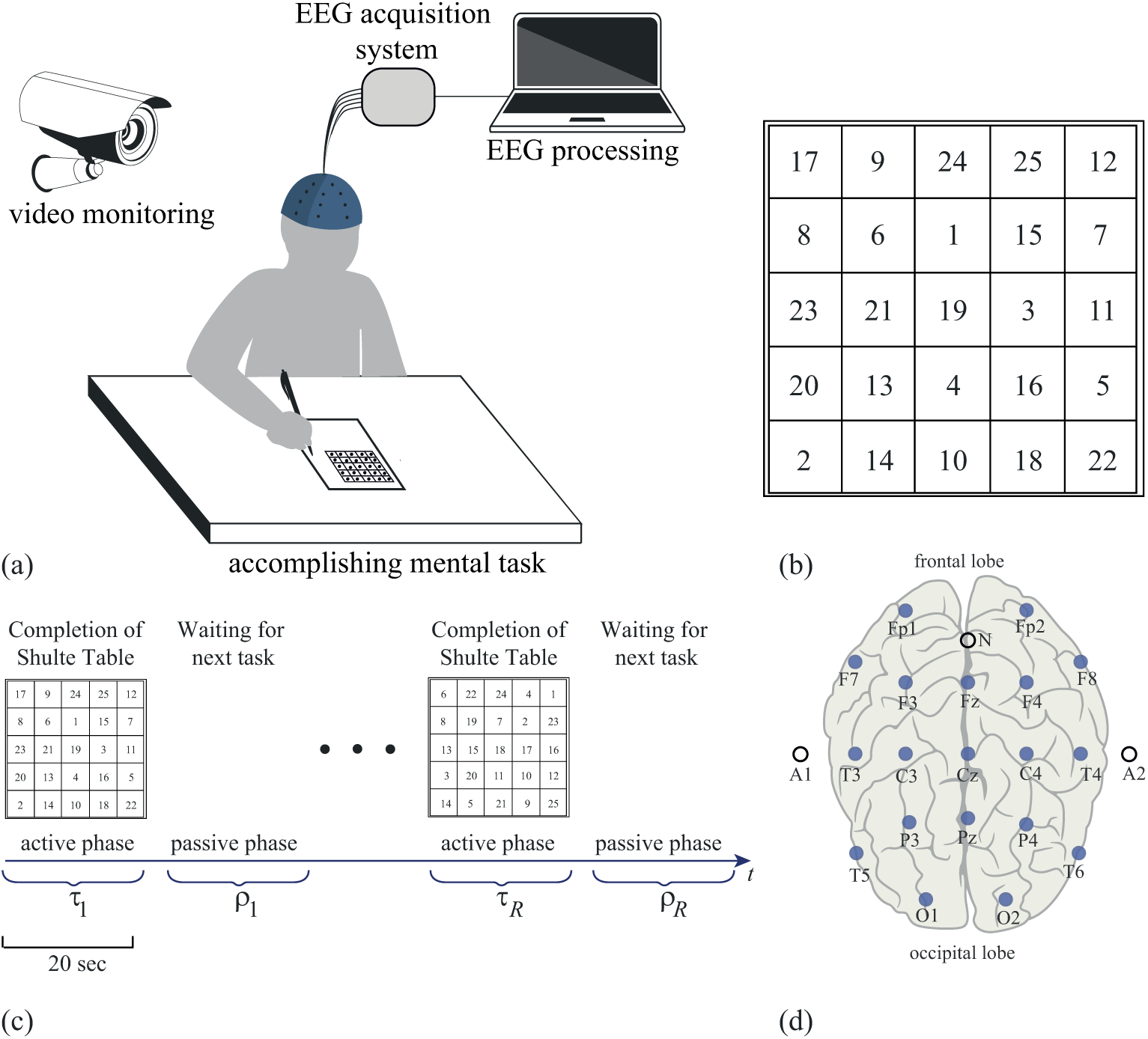
Experimental design. (a) Illustration of experimental procedure. (b) Typical Schulte 5 *×* 5 table. (c) Experimental design: completion of *R* Schulte tables (*i*-th active phase with length *τ_i_*) followed by *i*-th passive phase with length *ρ_i_* (waiting and preparing for next task). (d) Layout of EEG electrodes arranged according to standard international 10–20 system.

Simultaneously, the EEG signals of the brain activity were recorded. The multi-channel EEG data were acquired by using the amplifier BE Plus LTM manufactured by the EB Neuro S.P.A., Italy (atwww.ebneuro.com). The data from 19 electrodes with two reference electrodes (A1 and A2) were recorded with a 8-kHz sampling rate using a standard monopolar method. Adhesive Ag/AgCl electrodes attached to a special pre-wired head cap were used. The ground electrode N was located above the forehead, while two reference electrodes A1 and A2 were located on the mastoids. The EEG signals were filtered by a band-pass filter with cut-off points at 1 Hz (HP) and 300 Hz (LP), and a 50-Hz Notch filter. During the experiment, a video was recorded to save the time intervals corresponding to the active and passive experimental phases.

### The analysis of psycho-diagnostic tests

The current analysis of the 16PF answered items was based on 15 personality scales: Warmth (reserved vs. warm), Emotional Stability (reactive vs. emotionally stable), Dominance (deferential vs. dominant), Liveliness (serious vs. lively), Rule-Consciousness (expedient vs. rule-conscious), Social Boldness (shy vs. socially bold), Sensitivity (utilitarian vs. sensitive), Vigilance (trusting vs. vigilant), Abstractness (grounded vs. abstracted), Privateness (forthright vs. private), Apprehension (self-assured vs. apprehensive), Openness to Change (traditional vs. open to change), Self-Reliance (group-oriented vs. self-reliant), Perfectionism (tolerates disorder vs. perfectionistic), and Tension (relaxed vs. tense). All these scales were estimated for each participant.

The Schulte tables are frequently used as a psychodiagnostic test for studying properties of human attention. This is one of the most objective methods to determine working effectiveness and ability, as well as resistance to external interference. The time *τ_i_* of the *i*-th table completion was used to evaluate three standard test personal criteria: (1) work efficiency **WE** (the arithmetic mean of the table completion time), (2) warming-up work indicator **WU** (the ratio of the working time for the first table to **WE**), and (3) psychological stability **PS** (the ability to sustain the operational activity for a long time). These criteria are described by the following formulas:

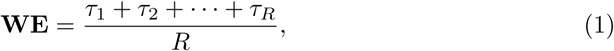

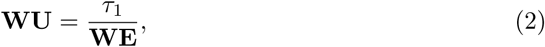

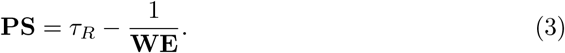

The work efficiency illustrates the attention consistency and performance. The resulted **WU** close to or lower than 1 indicates good warming-up, while 1 and higher means that the subject needs longer preparation time (warm-up) for the main work. The **PS** results close to 1 and less indicate a good psychological stability.

### EEG analysis

We analyzed the EEG signals recorded by 19 electrodes placed on the standard positions of the 10–20 international system [31] (see Fig. 1, (d)), using the continuous wavelet transform. The wavelet energy spectrum 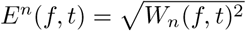 was calculated for each EEG channel *X_n_*(*t*) in the frequency range *f* ∈ [1, 40] Hz. Here, *W_n_*(*f, t*) is the complex-valued wavelet coefficients calculated as [32]

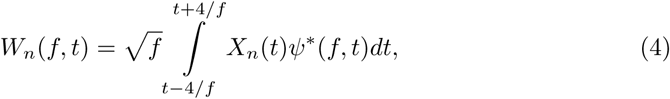

where *n* = 1*,…, N* is the EEG channel number (*N* = 19 being the total number of channels used for the analysis) and “*” defines the complex conjugation. The mother wavelet function *ψ*(*f, t*) is the Morlet wavelet often used for the analysis of neurophysiological data, defined as [32]

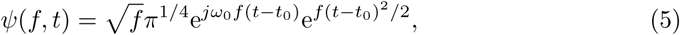

where *ω*_0_ = 2*π* is the central frequency of the mother Morlet wavelet.

Energy spectrum *E^n^*(*f, t*) was considered separately in the following frequency bands: delta (1–4 Hz), theta (4–8 Hz), alpha (8–13 Hz), beta–1 (13–23 Hz), beta–2 (24–34 Hz), and gamma (34–40 Hz) [33]. For these bands the values of wavelet energy 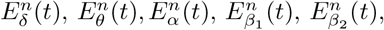 and 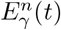 for each *n*-th EEG channel were calculated as

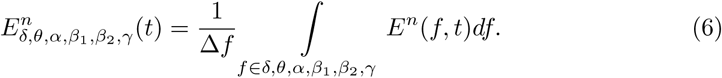

As a result, we considered the percentage of the spectral energy distributed in these bands, and calculated coefficients

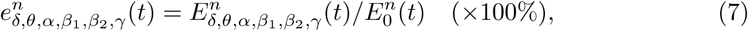

where *E*_0_(*t*) was defined as the whole energy and calculated as

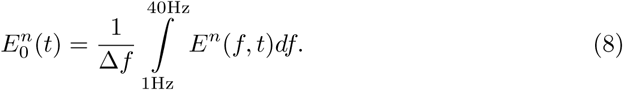

Finally, to describe the ratio between high frequency and low frequency brain activity for each channel, we introduced coefficient *ε^n^* defined as

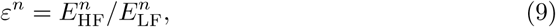
where

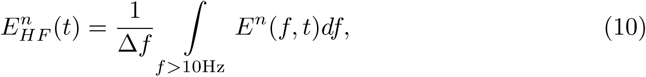

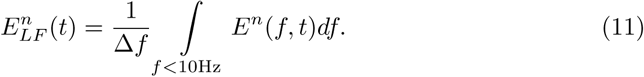

The coefficients *ε^n^* were calculated for each EEG channel for both the active and the passive phases. The obtained values of *ε_n_* were averaged over the channels located on the left and right hemispheres, defined respectively as

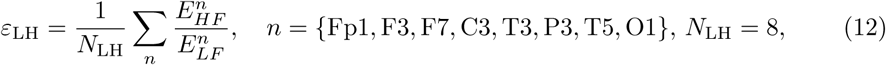

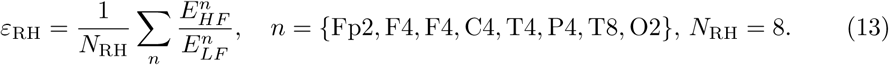

## Results

In the present work, we calculated the values of 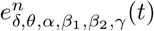 using Eq. (7)), for *n* = 1*,…* 19 EEG channels, which determined the percentage of the spectral energy belonging respectively to delta, theta, alpha, beta–1, beta–2, and gamma frequency bands, and characterized the degree of participation of the neural ensemble, located in the vicinity of the *n*-th recording electrode, in generation of the corresponding type of activity [7]. In Fig. 2 (b), we plot the values of 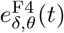 calculated for a single EEG trial recorded from the frontal lobe, specifically, from the F4 electrode. One can see, that when the active phase was replaced by the passive phase, the values of 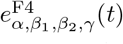 calculated for low frequencies (namely, delta, and theta frequency bands) rapidly increased, while the values of 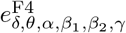, calculated for alpha, beta–1, beta–2, and gamma frequency bands, pronouncedly decreased. Such a dynamical behavior repeated itself during subsequent completions of the Schulte tables.

**Fig 2.**
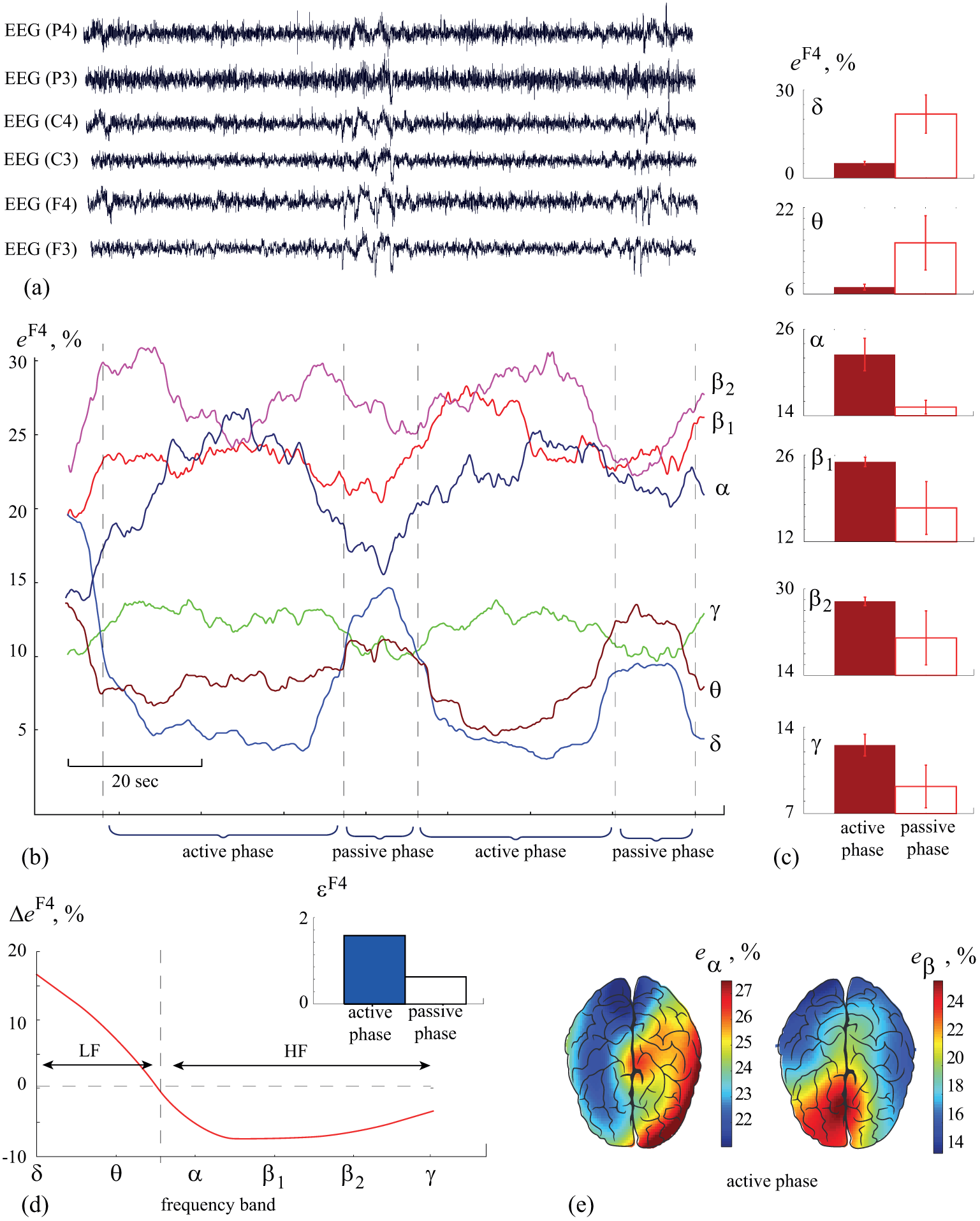
Quantification of EEG spectral properties. (a) typical EEG fragments, recorded by electrodes, arranged symmetrically in left and right hemispheres in frontal (F3, F4), central (C3, C4), and parietal (P3, P4) areas during active and passive phases. (b) Changes in spectral energy in different frequency bands. (c) Spectral energy averaged over *N* = 5 active and *N* = 5 passive phases (data are shown as mean ±SD). (d) Change in spectral energy during the transition from active to passive phase, calculated for each frequency band. *ε* defines the ratio between spectral energy in high (*f >* 10 Hz) and low (*f <* 10 Hz) frequency bands. (e) Typical distributions of spectral energy in alpha and beta–1 frequency bands during the active phase.

Figure 2 (c) shows the mean values of 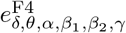 over the time intervals corresponding to *N* = 5 consecutive active and passive sessions. The distinctive features between the mean values obtained for the active and passive phases are displayed in Fig 2 (d), where the differences ∆*e*^F4^ between the mean values *e*^F4^ associated with the active and passive phases are plotted for each frequency band. One can see that in the low frequency range, which includes delta and theta frequency bands, such difference is positive (∆*e*^F4^ *>* 0), while in the high frequency range (alpha, beta–1, beta–2, and gamma frequency bands) it is negative (∆*e*^F4^ *<* 0).

According to this result, one can easily distinguish active and passive phases, based on the consideration of EEG properties, i.e., by comparing the energy of the spectral components belonging either to high (HF) or low (LF) frequency bands. For this purpose, it is convenient to use coefficient *ε^n^* (Eq. 9), which reflects the ratio between the values of spectral energy in the high and low frequency ranges. In particular, for the considered F4 electrode, the values of *ε*^F4^, shown in the inset histogram in Fig. 2 (d)) are significantly lower during the passive phase than during the active phase.

Thus, the time frequency analysis performed for a single EEG recording demonstrates a pronounced change in the ratio between the energy of high and low spectral components. At the same time, along with the features of time-frequency structure revealed in a single EEG, the spatio-temporal features of electrical brain activity also play an important role. This is mostly reflected in hemispheric differences commonly observed in electrical activity of the brain associated with the completion of mental tasks [34]. For instance, having considered the energy value calculated in alpha and beta–1 frequency bands during the active phase, one can see that these types of activity are localized in opposite hemispheres. Figure 2 (e) shows typical distributions of the spectral energy during the active phase in alpha and beta–1 frequency bands. It should be noted, that such a behavior of alpha activity is typical for arithmetic and visual-spatial tasks [35].

In order to understand this asymmetry, we considered coefficients *ε*_LH_ (12) and *ε*_RH_ (13), which were obtained by averaging coefficients *ε* calculated for EEG channels, belonging to the right and left hemispheres, respectively. In Fig 3 (a), the values *ε*_RH_ and *ε*_LH_ are shown for each of the 20 participants in the active (closed dots) and passive (open dots) phases. Having considered the obtained values, especially the difference between the ones calculated for active and passive phases, one can see that the subjects can be promptly divided into three groups. In Fig. 3 (a), the values *ε*_RH_ and *ε*_LH_ are shown for each group by different symbols. In group I (circles), coefficients *ε*_RH_ and *ε*_LH_ have practically the same values during the active and passive phases. In group II (squares), the active phase is associated with an increase in high-frequency activity in the right hemisphere and the passive phase with an increase in high-frequency activity in the left hemisphere. In group III (triangles), the transition from the active to the passive phase is associated with a pronounced increase in *ε*_RH_ and a decrease in *ε*_LH_.

**Fig 3.**
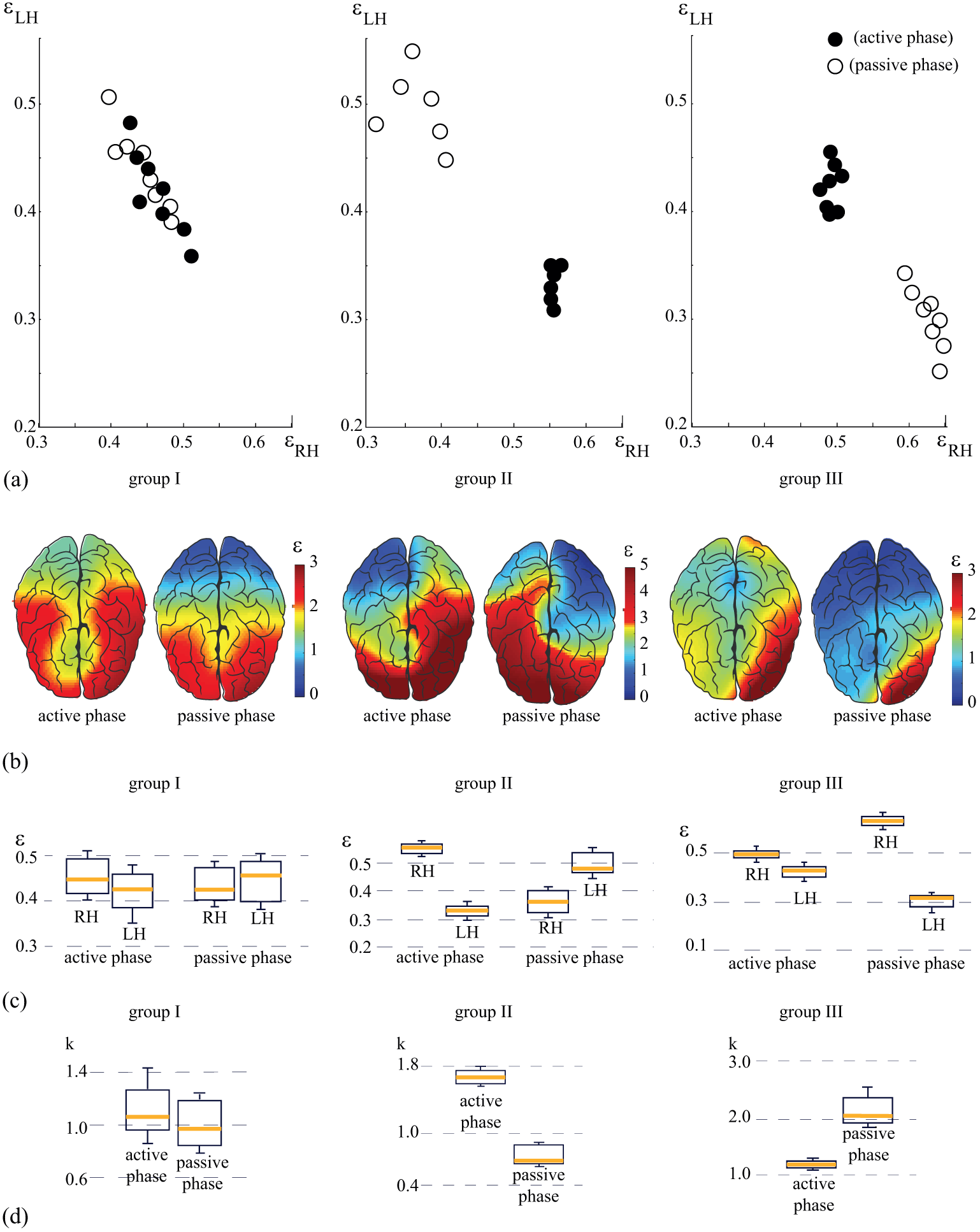
Three scenarios of cognitive activity during mental tasks processing. (a) Relation between energy of high-and low-frequency spectral components in the left (*ε*_LH_) and right (*ε*_RH_) hemispheres, calculated for active (closed dots) and passive (open dots) experimental phases. The distributions are shown for three subjects, each belonging to a particular group. (b) Coefficient *ε* showing the relation between energies of high-and low-frequency spectral components, calculated for each EEG channel during active (left-hand columns) and passive (right-hand columns) phases. (c) The ratio *ε* between energies of high-and low-frequency spectral components calculated for EEG channels belonging to left (LH) and right (RH) hemispheres during active and passive phases: medians (yellow bars), 25–75 percentiles (box), and outlines (whiskers). (d) The ratio *k* between the values of *ε* calculated for left and right hemispheres during active and passive phases: medians (yellow bars), 25–75 percentiles (box), and outlines (whiskers). Groups I and III contain *n* = 8 subjects, group II contains *n* = 6 subjects.

In Fig. 3 (b), the spatio-temporal representations of the values *ε* are shown for active and passive phases for each of the three groups. One can see that for group I, the brain activity during the active phase is characterized by hemispheric symmetry. In the passive phase, although the spatio-temporal structure of brain activity changes, the hemispheric symmetry persists. In group II, the spatio-temporal structure is significantly different. One can observe hemispheric asymmetry in active and passive phases, with high-frequency activity dominance during the passive phase. In group III, the hemispheric symmetry is observed. However, the high-frequency activity prevails in the right hemisphere as clearly shown in Fig. 3 (b). Thus, the difference in the brain activity in active and passive phases reveals itself as a change in the symmetry caused by a decrease in *ε* in the left hemisphere during the transition from the active to the passive phase.

The distinctive features of brain activity during the active and passive phases, observed in these three groups, are shown in Fig. 3 (c). The horizontal yellow bars indicate the median of *ε* calculated for the left (LH) and right (RH) hemispheres during the active and passive phases. In group I, the values of *ε* remain practically the same for different hemispheres in both the active and passive phases. In group II, the active phase is characterized by a sharp increase in *ε* in the right hemisphere (*ε*_RH_ *>* 0.5 vs *ε*_LH_ *<* 0.35). In the passive phase, the dynamics is reversed, namely, an increase in *ε* is observed in the left hemisphere (*ε*_RH_ *<* 0.4 vs *ε*_LH_ *>* 0.45). Finally, in group III, during the active phase, *ε* in the right hemisphere is slightly higher than that in the left hemisphere (*ε*_RH_ *>* 0.45 vs *ε*_LH_ *<* 0.45). During the passive phase, such a difference becomes greater (*ε*_RH_ *>* 0.6 vs *ε*_LH_ *<* 0.35).

## Discussion

It is known that the completion of mental tasks is associated with changes in neural activity, which can be detected in the EEG power spectrum. The role of low-frequency **delta activity** in mental tasks was studied in [36], where the authors reported on increasing delta EEG activity during mental tasks, associated with enhancing attention. Later [37], a relation between delta-oscillations and the performance of mental tasks was also identified. On the other hand, earlier works [38, 39] highlighted an increase in **theta activity** during mental efforts. Recently, a change in the activity level in the low-frequency *θ*-band was used to evaluate the dynamics of mental workload [40]. The relation between **alpha activity** and the completion of mental tasks was demonstrated yet in 1984 by Osaka [35], who detected changes in the amplitude and location of the peak alpha frequency in the power spectrum. Later, a significant role of alpha activity in memory and cognitive processes was identified [41]. Changes in the energy of high-frequency brain rhythms are usually related to cognitive activity, in particular, mental task completion [42]. For instance, the account of **gamma activity** for classification of mental tasks improves the accuracy [43].

According to Fig. 3 (c), one can see that electrical brain activity in each group follows a particular scenario defined, on one hand, by the lateralization of the brain function, and on the other hand, by specific transitions between active and passive phases. In order to quantitatively describe the observed scenarios, we calculated *k* = *ε*_RH_*/ε*_LH_, which reflects a degree of hemispheric asymmetry. These values are plotted for each group in Fig. 3 (d). One can see that group I is characterized by hemispheric symmetry in active and passive phases, which remains unchanged during active-passive phase transition (∆*k ≈* 0), where ∆*k* = *k*_passive_ – *k*_active_. For other groups, asymmetry and transition are observed between active and passive phases, and plotted in terms of *k* which can be described as ∆*k <* 0 and ∆*k >* 0, respectively.

The participants belonging to each of the three groups were subjected to psycho-diagnostic tests (see Methods). As a result, the values of **WE**, **WU**, and **PS**, which define the average time of task completion, average performance, and attention preservation, respectively, were estimated for each subject. In addition, the personality of each subject was described on the basis of The Sixteen Personality Factor Questionnaire. According to the results of the psycho-diagnostic test, the subjects were divided into three groups, which matched the classification obtained with the EEG analysis based on the psychological description of the Schulte tables performance.

The subjects from group I demonstrated bilateral EEG activity in both hemispheres during the Schulte tables tests. Simultaneously these subjects demonstrated a medium-low efficiency when performing the task. For them, the average time of the task completion was **WE** = 40.2 seconds, the average performance was **WU** = 1.07 (the target value was 1), and attention preservation was **PS** = 0.97 (the target value was 1). The subjects from this group could immediately perform unknown tasks and maintain their working efficiency at a relatively high rate, above a medium-low level. The psychological decryption of the tests included the remarks about the creativity in the test performance and fast switches to new tasks. In the personal test, such subjects had a pronounced tendency to work alone, high intellect, analytical mind, critical thinking, intolerance to uncertainty, and a delay in decision making. Moreover, they exhibited self-control, a lack of anxiety, a pronounced leadership, and a desire to dominate in the group. We hypothesize that the creativity and the attempt to optimize their work led to a decrease of their working efficiency.

The subjects in group II tried to develop a strategy to simplify the task performance. During the accomplishment of the first task, a maximum lateralization of high-frequency activity was present, i.e., the activity in the right hemisphere was much more pronounced. This means that during the first task, the strategy was not yet developed. During the next tasks, the burden in the right hemisphere in these subjects was reduced. As a result, the subjects from group II demonstrated higher working efficiency than the subjects from group I. The average time of task completion was **WE** = 33.6 seconds, the attention preservation was **PS** = 0.86, and the average performance was **WU** = 1.07. These subjects needed little time for adaptation and did not tire, being capable to effectively maintain a high working efficiency for a long time. Their personal profiles harmoniously combined high scores in intellect, emotional maturity, and self-control.

Unlike group II, the subjects from group III accomplished the task without any attempts to develop a strategy to simplify it. This was confirmed by the psychological test. Their working efficiency remained high: the average time of the task completion was **WE** = 33 seconds, the attention preservation was **PS** = 0.9, and the average performance was **WU** = 1.24. We assume that the subjects from this group would have difficulties to maintain a good working efficiency for a long time. Their personal tests showed a pronounced preference to work alone, low self-control, intolerance to uncertainty, and a delay in decision-making, which can be manifested by anxiety. They also demonstrated high intellect, an analytical mind, critical thinking, and a spirit for experimentation.

Figure 4 illustrates the correlation between the results obtained via the EEG study and the results of psycho-diagnostic tests and 16PF Questionnaire. The diagram In Fig. 4 (a) shows the results of the Cattell’s 16 Personality Factors Test for three groups. The data are displayed as the values of all primary factors of the 16PF Questionnaire, averaged over all subjects in each group. One can see that most of the factors have similar values in each group. At the same time, for some factors the corresponding values vary significantly from one group to another. Among these factors, one can distinguish Warmth (A), Reasoning (B), Emotional Stability (C), and Dominance (E), which are tabled and compared with the results of the EEG study and the psycho-diagnostic test in Fig. 4 (b).

**Fig 4.**
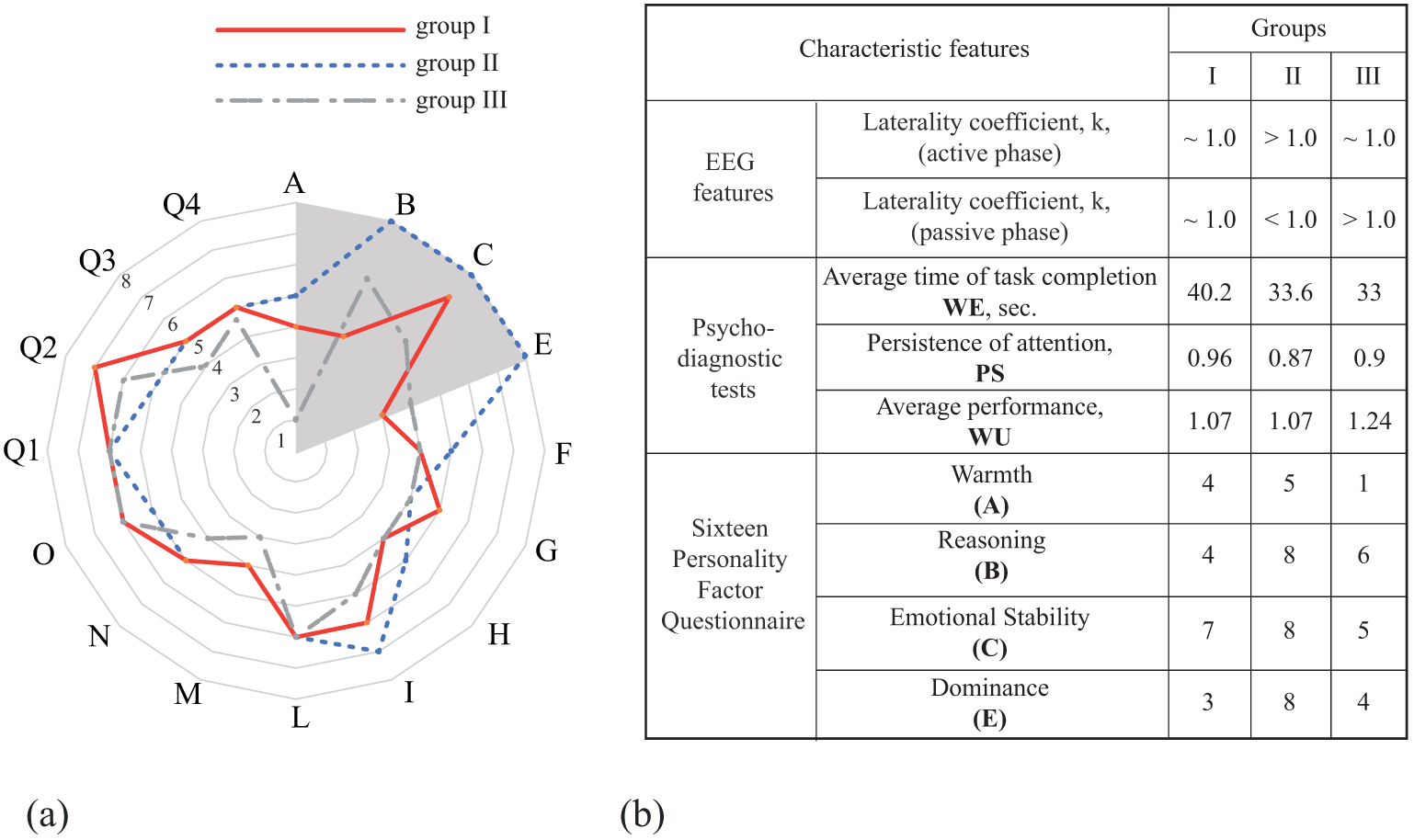
Sixteen personality factor questionnaire. (a) Values of primary factors of 16PF Questionnaire, averaged over subjects in each group: group I (dotted line), group II (solid line), and group III (dashed line). The dashed area highlights the factors for which significant changes between the groups are observed. (b) Correlation between results obtained via EEG and results of psycho-diagnostic tests and 16PF Questionnaire.

According to the results of personality classification based on the psycho-diagnostic test, one can see that different features of the EEG structure, i.e., lateralization and the ratio between energy of high-and low-frequency waves, reflects different personal qualities. It is important to note, that while EEG activity varied among different groups, it represented the same scenario inside each group. A similar behavior was observed in psychological classification, where three groups of subjects with similar personal profiles were identified.

Usually, majority of the scientific publications which aimed to reveal the EEG signatures of the cognitive activity describe the scenario, which is repeated from one subject to another. At the same time, we show that the differences occurred from one subject to another, can also be systematized. Different scenarios of cognitive activity can be identified among the subjects depending on the personality.

Our results confirm the hypothesis raised by Vingiano and William [44] about the existence of a relation between brain hemisphericity and personality. Our results are also in accordance with the work of [45], where the anxiety-related properties of personality, estimated via Cattel’s technique, were shown to correlate with the spectral power density (SPD) of EEG rhythms, in particular, beta–1 and beta–2. The authors claimed that the intense beta EEG rhythm correlates with highly situational and individual anxieties. At the same time, an individual’s emotional stability was found to be related to the alpha rhythm power.

Thus, the obtained results provide new knowledge in understanding the features of human personality by analyzing the relation between spatio-temporal and time-frequency EEG structure.

## Conclusion

We have analyzed the correlation between neurophysiological processes and personal characteristics during complicated mental tasks using a series of simple psycho-diagnostic tests to study human personality (the Schulte tables). To solve this task, we have considered spatio-temporal and time-frequency structures of multichannel EEGs in humans, who completed the Schulte tables. We have shown that EEG activity during the mental tasks varied from one subject to another. At the same time, three groups of subjects exhibiting similar features of neural activity were selected. The data of all subjects were independently analyzesd with the help of psycho-diagnostic tests in order to study their attention features and classify their personality profiles. As a result of this psychological classification, the subject were divided into three different groups. We have shown that the classification obtained via EEG study strongly correlated with the results of psycho-diagnostic tests. This, in turn, provided a possibility to characterize personality profiles based on the analysis EEG data.

We believe that our results can help in testing and diagnostic of personal skills and abilities to perform complex operational tasks. On the base of our findings, automatic intelligent systems can be developed to examine subject’s strong and weak points for high demanding purposes.

## Acknowledgments

This work has been supported by the Ministry of Education and Science of Russian Federation (Project RFMEFI57717X0282 of Russian Federal Target Programme).

